# TBC1D24 regulates mitochondria and endoplasmic reticulum-mitochondria contact sites

**DOI:** 10.1101/2024.09.19.613961

**Authors:** Sara Benhammouda, Justine Rousseau, Philippe M. Campeau, Marc Germain

## Abstract

The RabGAP protein TBC1D24 is mutated in several neurological disorders including DOORS syndrome (Deafness, Onycho-Osteodystrophy, Mental Retardation, Seizures). Although TBC1D24 comprises two highly conserved domains — the Tre2/Bub2/Cdc16 (TBC) domain and the TLDc domain — both of which seem to play critical roles in cellular functions, its exact physiological function within the cell remains poorly understood. Here, we show that TBC1D24 affects mitochondrial structure and activity. Specifically, both primary fibroblasts from patients with TBC1D24 mutations and cells in which TBC1D24 has been knocked down exhibit fragmented mitochondria, decreased ATP production, and reduced mitochondrial membrane potential. Importantly, loss or mutation of TBC1D24 also alters sites of contact between the endoplasmic reticulum (ER) and mitochondria (ERMCS). These sites are vital for mitochondrial fusion and fission processes, which regulate mitochondrial dynamics and, consequently, mitochondrial activity. Altogether, our results uncover a new role for TBC1D24 in the regulation of mitochondrial functions and ERMCS which likely contribute to the neurological dysfunction present in affected patients.

## Introduction

Mutations in the *TBC1D24* gene are associated with a range of neurological disorders, including various forms of epilepsy such as myoclonic epilepsy and drug-resistant epileptic encephalopathy, non-syndromic hearing loss, as well as DOORS syndrome (1-4). DOORS syndrome is a rare genetic disorder characterized by a combination of symptoms including Deafness, Onychodystrophy (abnormal growth of fingernails and toenails), Osteodystrophy (abnormal bone development), Retardation (intellectual disability), and Seizures (convulsions) (2) (5-7).

At the molecular level, the *TBC1D24* gene encodes a protein characterized by two highly conserved domains, each of which appears to play a critical role in cellular function. The TLDc (TBC/LysM domain-containing) domain is associated with cellular resistance to oxidative stress, although the underlying mechanisms remain poorly understood (8-10). This domain has been shown to interact with the V-ATPase (vacuolar-type H+-ATPase) complex and regulate its ability to acidify lysosomes (10-14). Interestingly, mutation in the *ATP6V1B2* gene that codes for a subunit of the V-ATPase have been associated with DOORS syndrome, suggesting a role for TBC1D24 in the regulation of the V-ATPase (13, 15, 16).

The second domain, the Tre2/Bub2/Cdc16 (TBC) domain, is integral to the regulation of vesicular transport. This domain is commonly found in a variety of Rab GTPase-activating proteins (Rab-GAPs), which are essential regulators of vesicular trafficking within cells (17-19). Consistent with this, TBC1D24 has been show to regulate Rab35 (20) and the small GTPase ARF6 (21-23), impacting neuronal growth and synaptic vesicle trafficking (21, 22, 24-26). Nevertheless, the physiological role of TBC1D24 remains poorly characterized (21, 22, 24-26). Interestingly, the Drosophila homologue of *TBC1D24* (Skywalker), has been found in a screen of proteins altering mitochondrial structure (27), suggesting the roles of TBC1D24 extend beyond endosomal alterations (20, 28). Consistent with this, symptoms of mitochondrial disease are also present in some TBC1D24-related epilepsy human patients (e.g., elevated plasma lactate, decreased mitochondrial respiratory chain complex activity), pointing to a potential involvement for TBC1D24 in mitochondrial function (29-32).

Given the pleiotropic roles of mitochondria in the regulation of cellular functions, including the activity of other organelles such as lysosomes (33-37), we investigated the effects of DOORS-related *TBC1D24* mutations on the structure and activity of mitochondria. Primary human fibroblasts from patients affected by DOORS with a mutation in *TBC1D24* showed reduced ATP levels and mitochondrial membrane potential that were associated with decreased mitochondrial length and connectivity. This was accompanied by alteration in endoplasmic reticulum (ER)-mitochondria contact sites (ERMCS). ERMCS play a key role in regulating mitochondrial structure and function by controlling the processes of mitochondrial fusion and fission (38-40). Altogether, our results indicate that TBC1D24 affects mitochondrial structure and function, and that this is associated with changes in ERMCS.

## Results

### Primary fibroblasts from DOORS patients have altered mitochondrial function

Mutations in TBC1D24 in humans lead to a number of neurological alterations, including DOORS syndrome. These mutations can occur in either of the two functional domains of the protein and are mostly observed as compound heterozygous. To better define the cellular roles of TBC1D24, we used a panel of primary fibroblasts from patients with *TBC1D24* mutations (Figure 1A). Most patient fibroblasts showed alterations in TBC1D24 protein expression (Fig 1B), with patient 2 (p.His487Glnfs^*^71; p.Cys105Arg) and 3 (p.His336GlnfsTer12^*^; c.1206+5G>A) fibroblasts having TBC1D24 below detection limit (Figure 1B), and thus acting as functional null mutants. As some conditions associated with *TBC1D24* mutations share phenotypic characteristics with mitochondrial diseases (30), and siRNA knockdown of the Drosophila homolog of *TBC1D24* leads to fragmented mitochondria (27), we then investigated the effect of these mutations on mitochondrial function in primary patient fibroblasts.

**Figure 1.**
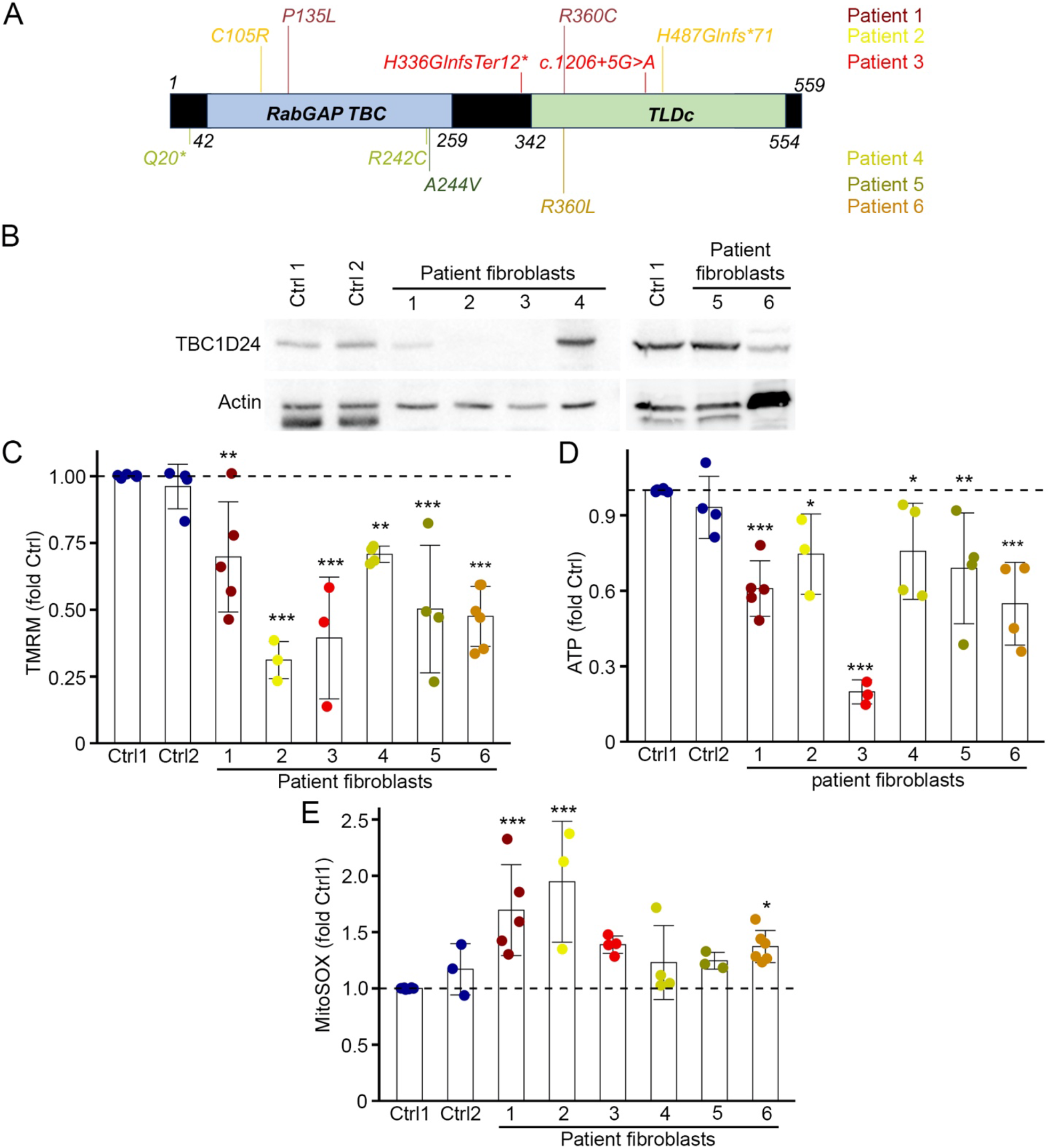
TBC1D24 mutations alter mitochondrial function. (A) Schematic representation of TBC1D24 with its two functional domains (RabGapTBC and TLDc) and the mutations present in the TBC1D24 patient fibroblasts used in this study. (B) Western blot showing TBC1D24 expression in the different control and patient lines. (C) Alterations in mitochondrial membrane potential in TBC1D24 mutant fibroblasts as measured using TMRM. Each point represents an individual experiment. Bars show the average ± SD. ^*^ p <0.05, ^**^ p<0.01, ^***^ p<0.001. One-way ANOVA. (D) ATP levels in TBC1D24 mutant fibroblasts. Each point represents an individual experiment. Bars show the average ± SD. ^*^ p <0.05, ^**^ p<0.01, ^***^ p<0.001. One-way ANOVA. (E) Mitochondrial ROS levels in DOORS syndrome fibroblasts as measured using MitoSOX. Each point represents an individual experiment. Bars show the average ± SD. ^*^ p <0.05, ^**^ p<0.01, ^***^ p<0.001. One-way ANOVA.

We first examined the effect of *TBC1D24* mutations on mitochondrial membrane potential (ΔΨ) using tetramethylrhodamine methylester (TMRM). All DOORS patients primary cell lines tested showed a decrease in mitochondrial ΔΨ, as compared to controls (Figure 1C). Consistent with this, total cellular ATP levels were also decreased in these cells (Figure 1D), suggesting that TBC1D24 mutations impair mitochondrial energy generation. As loss of mitochondrial function is often associated with an increase in reactive oxygen species (ROS), we also determined the amount of mitochondrial ROS in DOORS fibroblasts. Surprisingly, only some mutants (mutants 1, 2 and 6) had increased ROS as measured by MitoSOX (Figure 1E), suggesting that the decreased mitochondrial membrane potential and ATP levels were not directly associated with increased ROS.

### TBC1D24 mutations alter mitochondrial structure

Given that alterations in mitochondrial function often correlate with alterations in mitochondrial structure, our subsequent investigation aimed to assess the impact of TBC1D24 mutations on mitochondrial morphology and organization. To achieve this, we labeled mitochondria using an antibody targeting the outer mitochondrial membrane protein TOM20 and subsequently visualized them using confocal microscopy. Mitochondria were shorter or even fragmented in most mutant lines compared to control lines (Figure 2A-B). Consistent with this, mitochondrial connectivity, a measure of the branching of mitochondrial networks (41), was significantly decreased in mutants 1, 2 and 3 (Figure 2C). In contrast, two of the mutants (5 and 6) had increased mitochondrial connectivity but minimal changes in mitochondrial length (Figure 2B-C) despite decreased membrane potential and ATP levels (Figure 1C-D). Thus, while the exact changes in mitochondrial structure observed in TBC1D24 mutants depends on the nature of the mutation, all TBC1D24 mutants associated with DOORS syndrome have altered mitochondrial structure.

**Figure 2.**
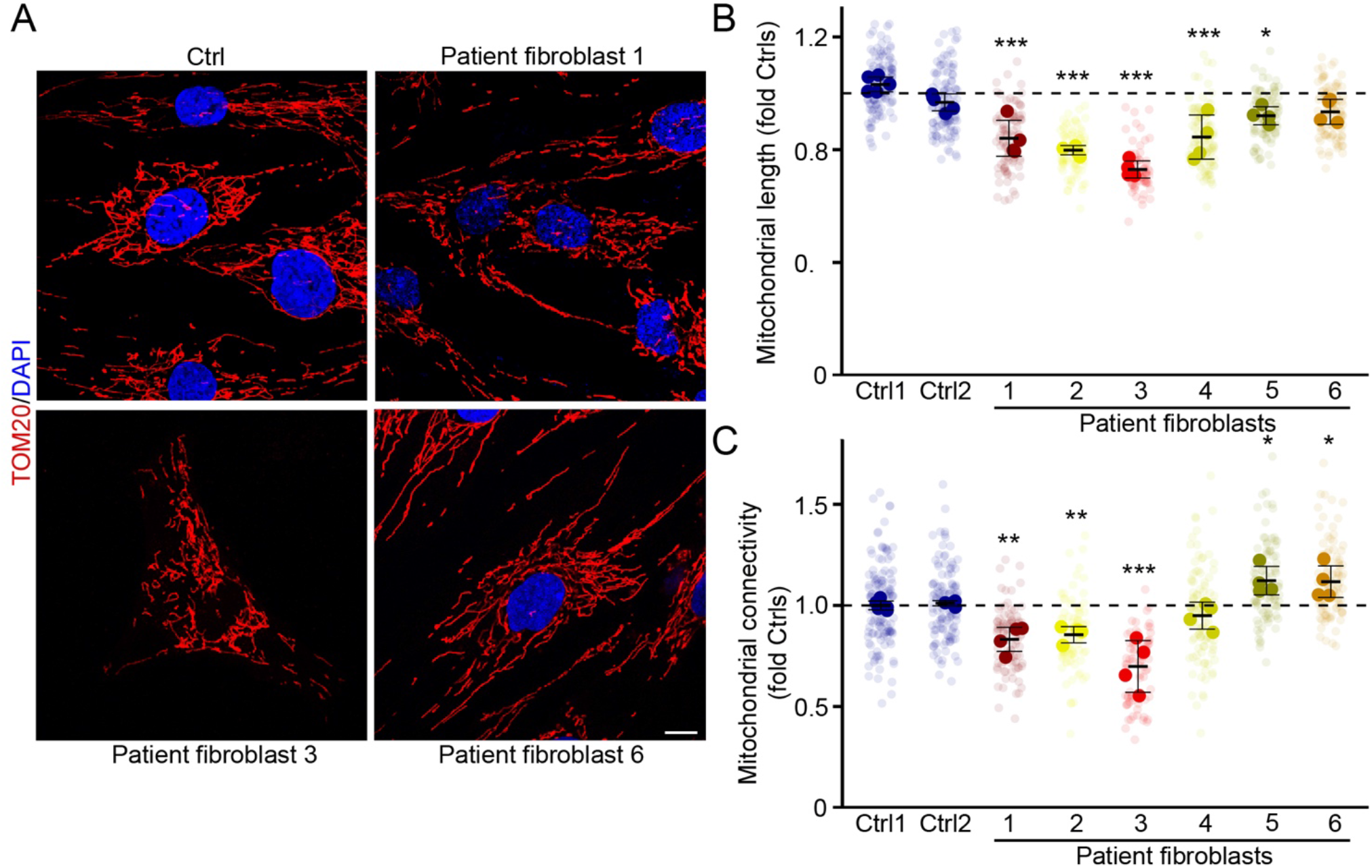
TBC1D24 mutations affect mitochondrial structure. (A) Representative confocal microscopy images of mitochondria (TOM20, Red) and nucleus (DAPI, Blue) of control and patient fibroblasts. Scale bar 10 µm. (B-C) Quantification of mitochondrial length (B) and connectivity (C) in the indicated control and patient fibroblasts imaged as in (A). Each point represents an individual experiment, with faded points representing individual cells within each experiment. Values were normalised to the average of controls. Bars show the average **±** SD. ^*^ p <0.05, ^**^ p<0.01, ^***^ p<0.001. One-way ANOVA.

### Downregulation of TBC1D24 in mouse embryonic fibroblasts recapitulates the mitochondrial alterations present in patient fibroblasts

Our findings indicate that patient fibroblasts have altered mitochondrial structure and function (Figure 1-2). However, as patients have distinct mutations and diverse genetic backgrounds, it is difficult to disentangle the contribution of the mutations and the genetic background. Thus, we next analyzed mitochondrial structure in mouse fibroblasts (MEFs) in which TBC1D24 was knocked down using siRNA (Figure 3A). Mitochondria were labelled with TOM20 and analysed by confocal microscopy (Figure 3B). As with patient fibroblasts, knockdown of TBC1D24 (siTBC) resulted in shorter mitochondria that were less connected (Figure 3C-D). We then determined if TBC1D24 knockdown affected mitochondrial activity. When compared to control siRNA, siTBC MEFs showed decreased cellular ATP levels and reduced mitochondrial membrane potential (Figure 3 E-F), consistent with patient fibroblasts. Nevertheless, siTBC MEFs displayed similar ROS levels as siCtrl cells (Figure 3G), consistent with the variable ROS levels observed in patient fibroblasts. Altogether, our results indicate that mutation or loss of TBC1D24 is associated with impaired mitochondrial structure and activity.

**Figure 3.**
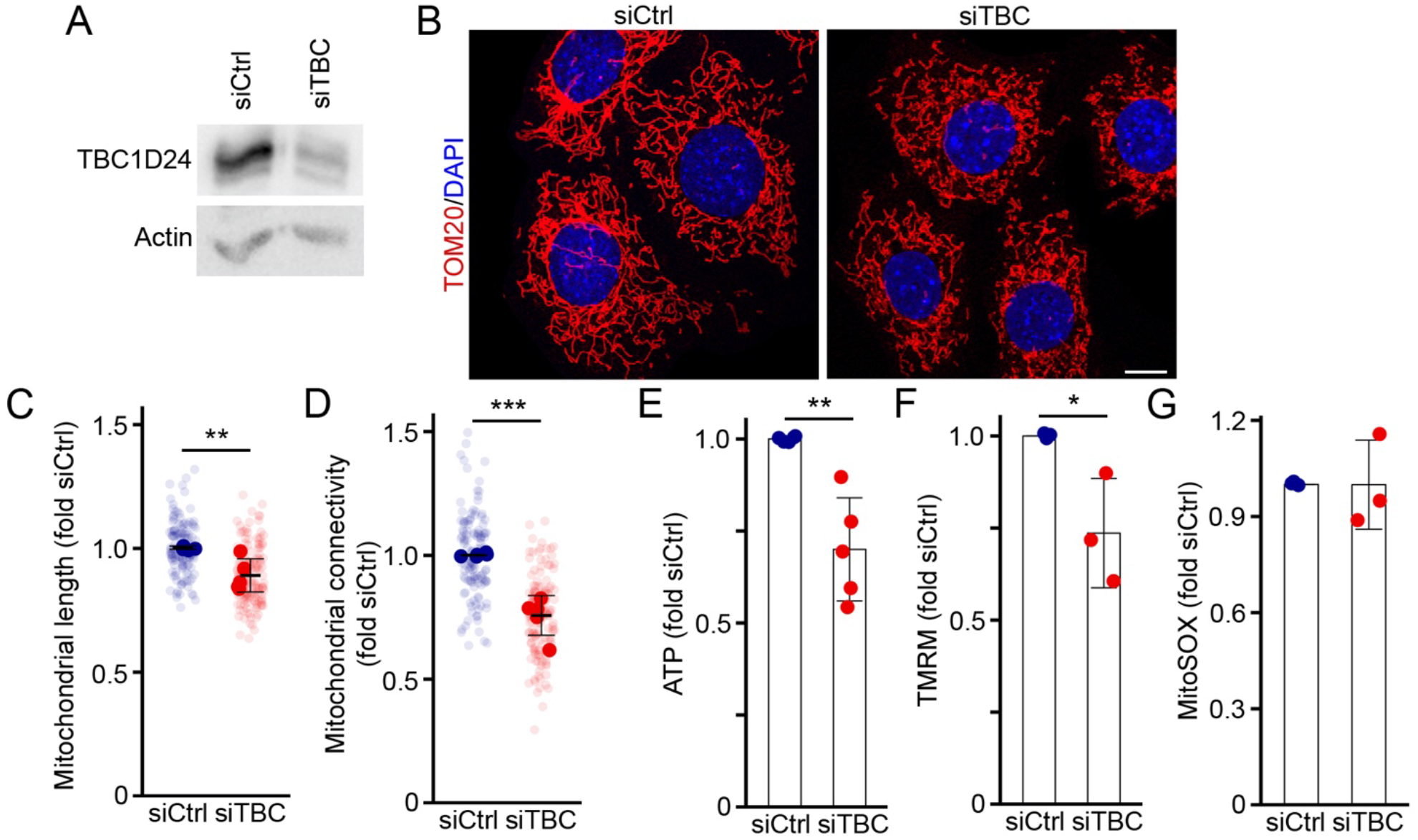
TBC1D24 knockdown causes mitochondrial fragmentation and impairs mitochondrial function. (A) Western blot showing the knockdown of TBC1D24 (siTBC) in MEFs (B) Representative confocal microscopy images of siCtrl and siTBC MEFs stained for mitochondria (TOM20) and nucleus (DAPI). Scale bars 10 µm. (C-D) Quantification of mitochondrial length (C) and connectivity (D) in siCtrl and siTBC MEFs. Each point represents an individual experiment, with faded points representing individual cells within each experiment. Bars show the average ± SD. ^**^ p<0.01, ^***^ p<0.001. Two-sided t-test. (E-G) Membrane potential (TMRM, E), ATP levels (F) and mitochondrial ROS (MitoSOX, G) in siCtrl and siTBC MEFs Each point represents an individual experiment. Bars show the average ± SD. ^*^ p <0.05, ^**^ p<0.01. Two-sided t-test.

### TBC1D24 expression affects mitochondrial structure

Having established a role for TBC1D24 in mitochondrial activity, we next determined whether increasing TBC1D24 expression could rescue the mitochondrial abnormalities observed in TBC1D24-deficient cells. For this, we first transfected different concentrations of a Flag-tagged version of wild type TBC1D24 (Flag-TBC1D24) in MEFs to determine the conditions where the exogenous protein is expressed at similar levels to the endogenous protein (Figure 4A). As a control, we also assessed mitochondrial structure in the transfected cell. Surprisingly, expression of a relatively large amount of Flag-TBC1D24 (1 µg) decreased mitochondrial length and connectivity (Figure 4B-C) similar to its downregulation, suggesting that TBC1D24 levels need to be finely tuned to maintain proper mitochondrial structure. However, transfection of a lower amount of Flag-TBC1D24 that does not affect mitochondrial length or connectivity in WT MEFs (0.3 µg) rescued mitochondrial structure in siTBC cells (Figure 4D-E), indicating that the defects observed in siTBC cells are TBC1D24-dependent.

**Figure 4.**
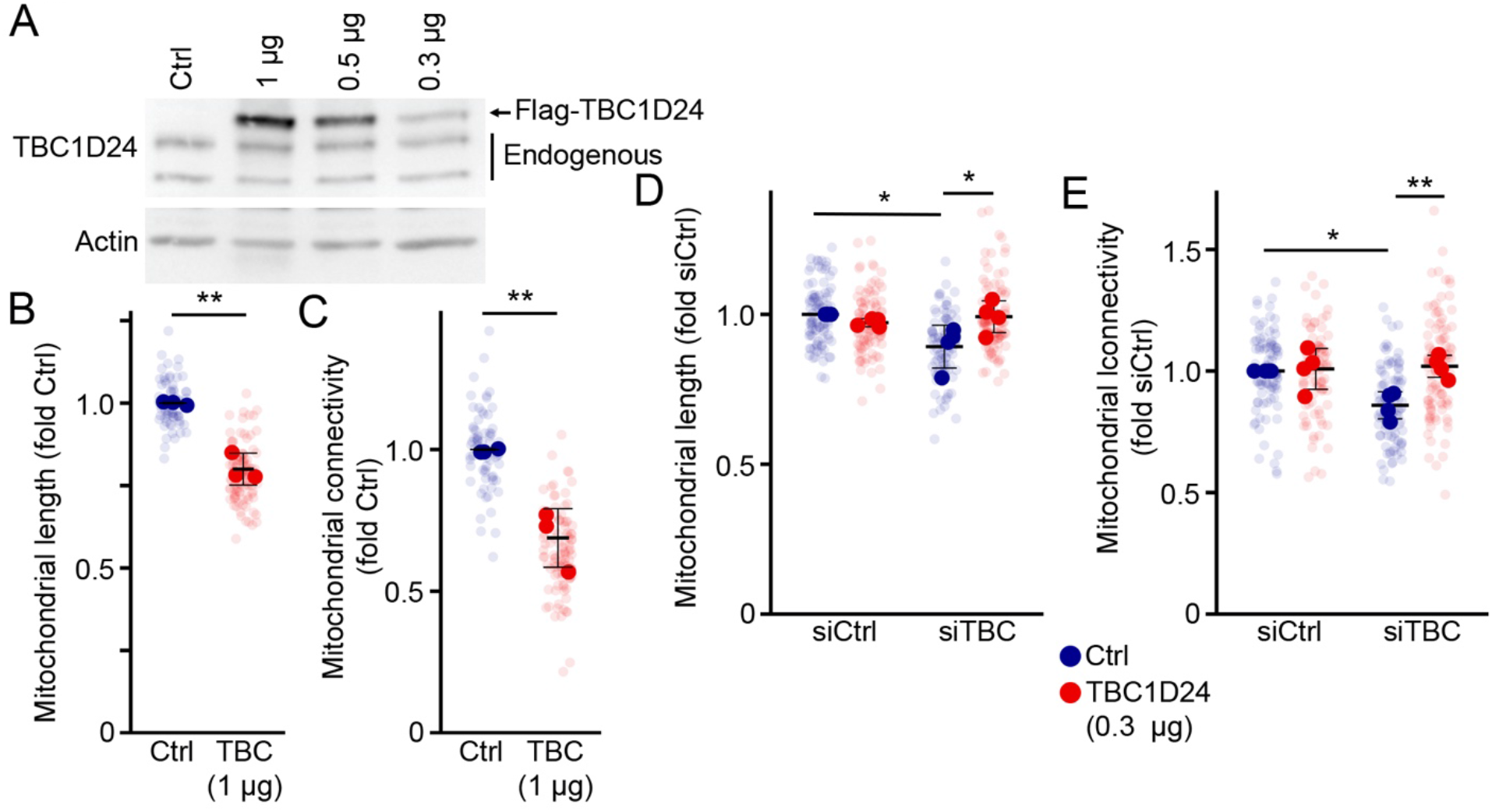
High levels of TBC1D24 promote mitochondrial fragmentation. (A) Western blot showing the expression of different amounts of Flag-TBC1D24 in MEFs. The top band is the transfected Flag-tagged protein. (B-C) Quantification of mitochondrial length (B) and connectivity (C) in MEFs expressing high levels (1 µg/well) of Flag-TBC1D24 (TBC). Each point represents an individual experiment, with faded points representing individual cells within each experiment. Bars show the average ± SD. ^**^ p<0.01. Two-sided t-test. (D-E) Rescue of mitochondrial length (D) and connectivity (E) in siTBC MEFs expressing low levels of Flag-TBC1D24 (0.3 µg/well). Each point represents an individual experiment. Bars show the average ± SD. ^*^ p<0.05, ^**^ p<0.01. Two-way ANOVA.

### TBC1D24 mutations alters the structure of the endoplasmic reticulum (ER)

Since TBC1D24 mutations associated with DOORS syndrome cause alterations in mitochondrial structure and function, we determined whether it localises to mitochondria by expressing Flag-TBC1D24 in control human fibroblasts. TBC1D24 did not colocalize with mitochondria (Figure 5A), suggesting that the effect on mitochondria might be indirect. As the endoplasmic reticulum (ER) plays an important role in the regulation of mitochondrial structure and function through the generation of ER-mitochondria contact sites (ERMCS), we also determined whether TBC1D24 localizes to the ER. Flag-TBC1D24 did indeed partially colocalize with the ER marker RTN4 (Figure 5B), from where it could potentially affect mitochondria.

**Figure 5.**
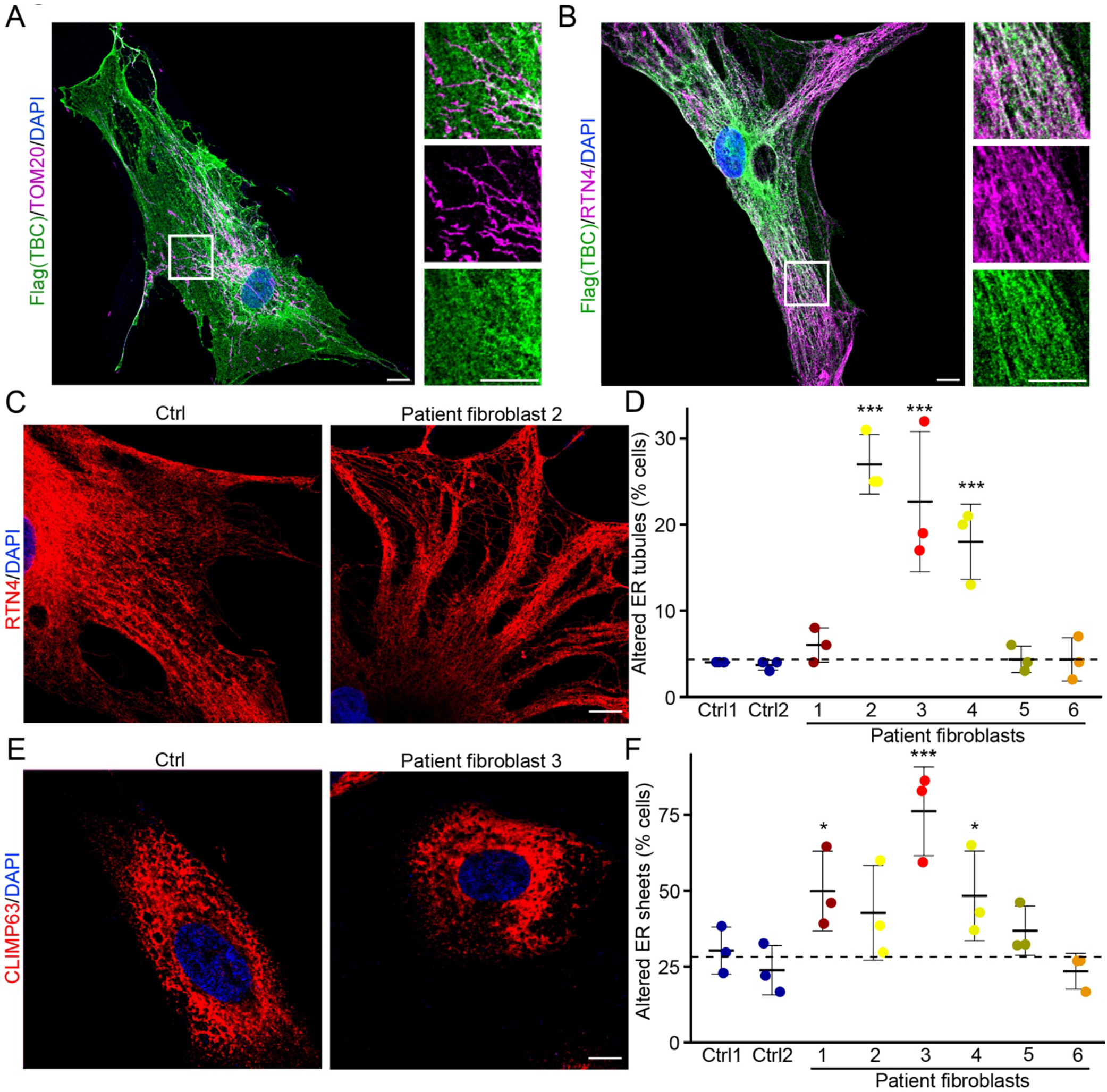
TBC1D24 mutations affect the structure of the endoplasmic reticulum (ER). (A-B) Representative images showing the localisation of Flag-TBC1D24 (Flag, Green) in transfected primary fibroblasts, along with mitochondria (TOM20, Magenta)(A) or ER tubules (RTN4, Magenta)(B). (C-D) TBC1D24 mutations affect the structure of ER tubules. (C) Representative images ER tubules (RTN4, Red) and nuclei (Dapi, Blue) in control and patient fibroblasts. Scale bars 10 µm (D) Quantification of ER tubule phenotypes from images in (C). Cells with ER tubules similar to the patient fibroblast shown in (C) were considered as altered. Each point represents an individual experiment. Bars show the average **±** SD. ^***^ P<0.001. One-way ANOVA. (E-F) TBC1D24 mutations affect the structure of ER sheets. (E) Representative images ER tubules (CLIMP63, Red) and nuclei (Dapi, Blue) in control and patient fibroblasts. Scale bars 10 µm (F) Quantification of ER sheets phenotypes from images in (E). Cells with ER tubules similar to the patient fibroblasts shown in (E) were considered as altered. Each point represents an individual experiment. Bars show the average **±** SD. ^*^ p<0.05, ^***^ P<0.001. One-way ANOVA.

We thus determined whether the ER structure was affected in TBC1D24 patient cells. The ER is composed of two domains that are connected but physically and functionally distinct: ER tubules at the periphery and ER sheets surrounding the nucleus (42, 43). We first investigated the structure of ER tubules using RTN4 as a marker (44). In contrast to control cells where tubular ER extended as a web-like network, a large fraction of mutant cells showed altered structure with coiled tubules and little ER between them (Figure 5C). Similar to the alterations in mitochondrial structure, coiled ER tubules were apparent in patient fibroblasts 2, 3 and 4, but not in patient fibroblasts 5-6. (Figure 5D). We then assessed changes in ER sheets using CLIMP63 as an ER sheet marker (45). Patient fibroblasts 1, 3 and 4 had altered ER sheets that appeared thicker and flatter than their control counterpart (Figure 5E, F). Altogether, our results indicate that TBC1D24 mutations are associated with alterations in both mitochondria and the ER.

### TBC1D24 mutation affects ER-mitochondria contact sites

ERMCS control mitochondrial dynamics and the transfer of material including calcium and lipids between the two organelles (46, 47). Given that TBC1D24 mutations affect the structure of mitochondria and the ER, we determined if TBC1D24 mutations also affect ERMCS. To address this question, we used a proximity ligation assay (PLA) for Calnexin (an ER marker) and TOM20 (mitochondrial marker) to detect ERMCS. To evaluate the specificity of the interaction, we co-labeled the cells for Calnexin and TOM20. When the Calnexin and TOM20 signals overlapped, clear PLA foci were detected (Figure 6A), indicating the presence of ER-mitochondria contact sites. Three TBC1D24 mutants (2, 3 and 6) showed a significant increase in PLA foci (Figure 6A-B), indicating an increase in ER-mitochondrial interaction sites. Interestingly, in contrast to mutants 2 and 3, mutant 6 exhibited normal mitochondria and ER phenotypes (Figures 2, 5), suggesting that ERMCS alterations are not directly linked to mitochondrial fragmentation. Nevertheless, siTBC MEFs also showed a significant increase in TOM20-Calnexin-positive ERMCS compared to siCtrl cells (Figure 6C-D). Altogether, these results suggest that TBC1D24 plays a role in the control of contact sites between mitochondria and the ER.

**Figure 6.**
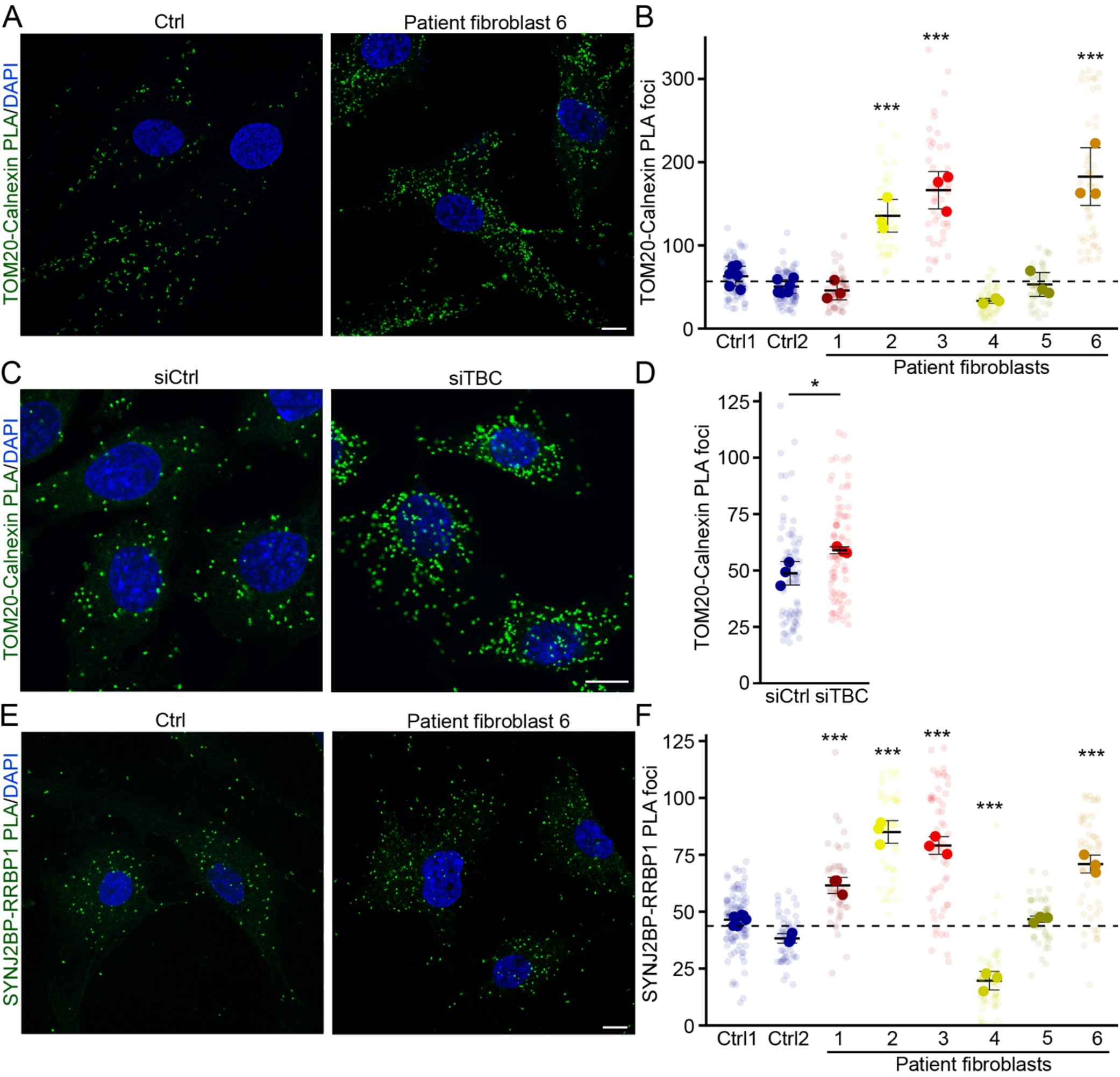
TBC1D24 mutations increase ER-mitochondria contact sites. (A-B) TBC1D24 mutations increase TOM20-Calnexin PLA foci. Representative images of control and TBC1D24 patient fibroblasts showing PLA foci for the ER protein Calnexin and the mitochondrial protein TOM20 (Green), along with nuclei (DAPI, Blue). The number of PLA foci is quantified in (B) Each point represents an individual experiment, with faded points representing individual cells within each experiment. Bars show the average ± SD. ^***^ p<0.001. One-way ANOVA. (C-D) TBC1D24 mutations increase SYNJ2BP-RRBP1 PLA foci. Representative images of control and TBC1D24 patient fibroblasts showing PLA foci for the ER protein RRBP1 and the mitochondrial protein SYNJ2BP (Green), along with nuclei (DAPI, Blue). The number of PLA foci is quantified in (B) Each point represents an individual experiment, with faded points representing individual cells within each experiment. Bars show the average ± SD. ^***^ p<0.001. One-way ANOVA. (E-F) Knockdown of TBC1D24 in MEFs increase TOM20-Calnexin PLA foci. Representative images of siCtrl and siTBC MEFs showing PLA foci for the ER protein Calnexin and the mitochondrial protein TOM20 (Green), along with nuclei (DAPI, Blue). The number of PLA foci is quantified in (B) Each point represents an individual experiment, with faded points representing individual cells within each experiment. Bars show the average ± SD. ^*^ p<0.05. Two-sided t-test

TOM20 and Calnexin are general markers for their specific organelle that are not necessairly direct binding partners in ERMCS. Thus, to further validate our findings, we used validated ERMCS interacting partners, the mitochondrial protein SYNJ2BP and the ER sheet marker RRPB1, to measure ERMCS by PLA (48, 49). Similar to the TOM20-Calnexin pair, SYNJ2BP-RRBP1 PLA foci were increased in mutants 2, 3 and 6 (Figure 6E-F). Interestingly, ERMCS contacts were also increased in mutant 1, while they were decreased in mutant 4 (Figure 6E-F). Altogether, our results suggest that TBC1D24 participate in the regulation of the contact sites between mitochondria and the ER, which could explain the mitochondrial phenotype observed in these cells.

## Discussion

Mutations in the TBC1D24 gene are associated with a range of neurological disorders, underscoring the gene’s essential role in neuronal function. The encoded protein contains conserved TBC and TLDc domains, whose specific roles and interactions are not yet fully understood (1-4). Consistent with the known role of TBC domains in the regulation of Rab GTPases, previous reports have indicated that TBC1D24 plays an important role in synaptic vesicle trafficking, notably their recycling (20-22, 24, 28). Here, we demonstrate that TBC1D24 also modulates mitochondrial structure and function. Specifically, we demonstrate substantial mitochondrial dysfunction in TBC1D24 fibroblasts derived from patients with DOORS syndrome. This dysfunction is evidenced by decreased ATP levels, reduced mitochondrial membrane potential, and altered mitochondrial morphology, indicating that TBC1D24 mutations may have a profound impact on these cellular processes.

Mitochondrial dysfunction is intimately connected to neurodegeneration through several mechanisms. Neurons rely heavily on mitochondria for ATP production, making them particularly vulnerable to impaired mitochondrial ATP synthesis. Dysfunctional mitochondria further exacerbate this vulnerability by generating excessive amounts of reactive oxygen species (ROS) (50-54). DOORS syndrome symptoms (hearing and/or vision problems, learning disabilities, developmental delay, intellectual disability, epilepsy) are also associated with mitochondrial dysfunction and some patients affected by DOORS syndrome with TBC1D24 mutations exhibit reduced activity of mitochondrial respiratory chain (29, 30). The role of TBC1D24 in regulating mitochondria that we uncovered is thus consistent with this. This is also supported by the observation that knockdown of Skywalker, the Drosophila homologue of TBC1D24, also causes mitochondrial fragmentation, although the functional consequences were not further investigated in that study (27).

We also found that mutations in TBC1D24 disrupt ER structure and affect ERMCS. While ERMCS regulate mitochondrial dynamics, they also affect other processes including protein folding, calcium balance, and lipid transport (47, 55). In this context, it is noteworthy that while most TBC1D24 mutations tested affect ERMCS, some did not show a corresponding increase in mitochondrial fragmentation. This suggests that some of the roles of TBC1D24 related to ERMCS are independent of mitochondrial dynamics. This complexity may help explaining the diverse clinical manifestations observed in DOORS syndrome.

The multiple effects of TBC1D24 loss of function on cellular activities (including ERMCS, mitochondrial function, synaptic vesicle trafficking and potential lysosomal defects through affecting the V-ATPase (10-14)) leads to the question of the mechanistic links between these alterations and their causal relationship to the disease. Previous studies have shown that mitochondrial dysfunction, as evidenced by altered energy production, calcium signaling, oxidative stress, and autophagy pathways, impacts ER function and contributes to ERMCS instability and lysosomal dysfunction (50, 56, 57). This could suggest that mitochondrial dysfunction is the primary defect present in TBC1D24 patient cells. However, primary defects in other organelles also indirectly affect mitochondria. For example, altered ERMCS can also compromise mitochondrial activity and impact the endocytic pathway by disrupting protein synthesis, folding, trafficking, altering membrane dynamics, vesicle formation, and impairing calcium-dependent vesicle fusion events (58-61). Similarly, disruption of the endocytic compartment, especially lysosomes, leads to the accumulation of dysfunctional mitochondria (62-64), highlighting the complex interplay of organelle dysfunction in cellular processes affected by TBC1D24 mutations.

A second possibility is that mutations in the TBC or TLDc domains lead to distinct phenotypes. However, we did not find a direct correlation between the location of mutations within the TBC or TLDc domains and the resulting cellular phenotypes. This analysis is however complicated by the fact that different TBC1D24 mutants are expressed at different levels. In fact, patient fibroblasts where TBC1D24 was not detected by western blot (2 and 3) generally had more severe phenotypes and knocking down TBC1D24 in MEFs recapitulated the phenotypes of patient fibroblasts. The phenotypic outcomes in cells with TBC1D24 mutations might thus be only partially or indirectly linked to specific mutations in either domain. In fact, cells seem to be very sensitive to changes in TBC1D24 expression levels as overexpressing it also alters mitochondrial structure. The variability we observed in patient fibroblasts is also in line with patients with TBC1D24 mutations exhibiting significant phenotypic variability. For example, while patients 1-4 from the present study developed multiple neurological issues including DOORS syndrome (1, 65, 66), patients 5 and 6 had only seizures (65, 67). Understanding these complexities is essential for unraveling the underlying pathophysiology of DOORS syndrome and developing personalized treatment strategies that consider both genetic factors and organelle-specific manifestations of the disease.

Our findings collectively highlight the important role of TBC1D24 not only for synaptic vesicle trafficking but also for the proper function of mitochondria and the ER. While this a novel mechanism by which TBC1D24 mutations can lead to DOORS syndrome and other neurological disorders, further research is required to fully uncover the physiological role of TBC1D24 protein and its involvement in metabolic pathways. Such investigations will offer a deeper comprehension of how mutations in TBC1D24 lead to neurological disorders.

## Methods

### Cell Culture

Primary human fibroblasts (controls and TBC1D24 patients) were generated from skin biopsies, collected as part of a research protocol approved by the CHU Sainte-Justine institutional review board, and written informed consent from participants was obtained. The cells were cultured in Dulbecco’s modified Eagle’s medium (DMEM) containing 10% fetal bovine serum (FBS), supplemented with Penicillin/Streptomycin (100 IU/mL/100 μg/mL). Immortalized Mouse Embryonic Fibroblasts (MEFs)(68) were cultured in DMEM supplemented with 10% fetal bovine serum.

### siRNA treatment

MEFs were seeded into 6-well plates and transfected with ON-TARGETplus Mouse Tbc1d24 siRNA (Dharmacon™ siRNA solutions, L-040513-01-0005) and negative siRNA (Thermo Fisher Scientific, Silencer Select, 4390843) using the siLenFect lipid reagent (Bio-Rad, 1703361). After 36 hours, the cells were collected for either western blot analysis or seeded onto coverslips for immunofluorescence studies.

### Immunofluorescence

Cells were seeded onto glass coverslips (Fisherbrand, 1254580) and allowed to adhere overnight. After fixation with 4% paraformaldehyde for 15 minutes at room temperature (RT), cells were permeabilized and blocked with PBS containing 0.1% Triton X-100 and 1% BSA for 10 minutes at room temperature. The cells were then incubated for 1 hour at room temperature in blocking buffer with the following primary antibodies: TOM20 (Rb, Abcam, ab186735, 1:250), CLIM63 (Mo, ENZO, ENZ-ABS669, 1: 200), RTN4 (Rb, Bio-Rad, AHP1799, 1:200), FLAG M2 (Mo, Sigma-Aldrich, F1804, 1:200). The cells were incubated with fluorescent tagged secondary antibody (Jackson Immunoresearch, 1:500) and DAPI (Invitrogen, Thermo Fisher, D1306, 1:100).

### Microscopy

Images were acquired using a Leica TSC SP8 confocal microscope equipped with a 63x / 1.40 oil objective, utilizing optimal resolution for the wavelengths used (as determined by Leica software).

### Image processing and analysis

All image manipulation and analysis were done in Image J/Fiji. For the analysis of mitochondrial length and connectivity, mitochondrial images taken with the confocal microscope were first segmented (converted into binary images) using ImageJ/Fiji, then processed with the Momito algorithm we developed (41). This software generates a skeleton representing the mitochondrial network and identifies the structural elements of this network (mitochondrial tubules, junctions, and ends). All possible connection models linking these structural elements were then determined and compiled to generate the overall distribution of the mitochondrial network.

### Proximity Ligation Assay (PLA)

PLA was as previously described (69). Briefly, Cells were cultured on glass coverslips and fixed with 4% paraformaldehyde. After antigen retrieval, cells were permeabilized with 1% BSA/0.2% Triton X-100 in PBS for 15 minutes at room temperature. Mitochondria were labeled with antibodies against TOM20 or SYNJ2BP (Rb, Sigma-Aldrich, HPA000866, 1:200), and the ER was labeled with antibodies against CALNEXIN (Mo, Millipore, MABF2067, 1:200) or RRBP1 (Mo, Thermo Fisher Scientific, MA5-18302, 1:200). PLA assays were conducted using the Duolink In Situ green kit Mouse/Rabbit (Sigma Aldrich) according to the manufacturer’s instructions. The primary antibodies were subsequently labeled with fluorescent-tagged secondary antibodies. Total PLA foci were manually counted at the sites of TOM20-CALNEXIN and SYNJ2BP-RRBP1 signal.

### ATP Assay/Luciferase activity assay

Cells were seeded in 96-well plates and, after 24 hours, the substrate solution from the Luciferase Assay System (Promega-CellTiter-Glo® 2.0 Cell Viability Assay-G9241) was added according to the manufacturer’s instructions. The plate was then incubated and kept in the dark for 10 minutes. Finally, luminescence was measured using a plate reader.

### Measurement of mitochondrial ROS production

Cells were stained with MitoSOX™ Mitochondrial Superoxide Indicator (Thermo-Fisher-MitoSOX™ Mitochondrial Superoxide Indicators-M36008) following the manufacturer’s protocol. Briefly, cells were incubated with 5 µM MitoSOX reagent at 37°C for 20 minutes, protected from light. Stained cells were then immediately analyzed using flow cytometry to assess mitochondrial superoxide production, ensuring minimal light exposure to prevent photobleaching and degradation of the fluorescent signal.

### Measurement of mitochondrial membrane potential

The mitochondrial membrane potential was measured using tetramethylrhodamine methyl ester (TMRM) from the MitoProbe™ TMRM Assay Kit (Thermo Scientific™). Cells were harvested and incubated with TMRM at 37°C for 20 minutes. Following incubation, the cells were washed with PBS, and fluorescence measurements were analyzed using flow cytometry.

### Western Blot

Cells were lysed in a buffer containing 10 mM Tris-HCl (pH 7.4), 0.5 mM EDTA, 150 mM NaCl, 1% Triton X-100, 50 mM sodium fluoride, and a Sigma-Aldrich protease inhibitor cocktail. Protein concentrations were determined using the BioRad DC protein assay kit. For SDS-PAGE, 25 μg of proteins were mixed with Laemmli buffer, separated by electrophoresis, transferred to nitrocellulose membranes, and probed with specific antibodies TBC1D24 (Mo, Santa Cruz, sc-515806, 1:1000), HRP-tagged Actin (HRP, Santa Cruz, sc-47778, 1:10,000). Membranes were incubated with horseradish peroxidase-conjugated secondary antibodies, visualized using enhanced chemiluminescence with a Bio-Rad imaging system.

### Statistical analysis

All graphs and statistical analysis were done using R. Immunofluorescence data was quantified from at least 3 experiments and images representative of at least three independent experiments are shown. Data is represented as the average ± SD as specified in figure legends. Statistical significance was determined using Student’s t test (between 2 groups), Dunnet test for the comparisons of individual control and patient fibroblasts against the average of control fibroblasts, or one-way ANOVA with a Tukey post hoc test (multiple comparisons).

## Acknowledgements

We thank the patient families and the collaborating physicians (authors in the cited manuscripts) for sharing patient fibroblasts after informed consent. This work was supported by an Acceleration grant from le Centre d’Excellence en Recherche sur les Maladies Orphelines-Fondation Courtois (CERMO-FC) to MG and PMC and a grant from the Natural Sciences and Engineering Research Council of Canada to MG. SB was the recipient of a CERMO-FC scholarship and holds a FRQS PhD scholarship.

